# Influenza A (N1-N9) and Influenza B (B/Vic and B/Yam) Neuraminidase Pseudotypes as Tools for Pandemic Preparedness and Improved Influenza Vaccine design

**DOI:** 10.1101/2021.09.19.460966

**Authors:** Kelly A.S. da Costa, Joanne Marie M. Del Rosario, Matteo Ferrari, Sneha Vishwanath, Benedikt Asbach, Rebecca Kinsley, Ralf Wagner, Jonathan L. Heeney, George W. Carnell, Nigel J. Temperton

**Affiliations:** Viral Pseudotype Unit, Medway School of Pharmacy, The Universities of Greenwich and Kent at Medway, Chatham, United Kingdom; Department of Physical Sciences and Mathematics, College of Arts and Sciences, University of the Philippines Manila, Manila, Philippines; DIOSynVax, Cambridge, United Kingdom; Department of Veterinary Medicine, University of Cambridge, Cambridge, United Kingdom; Institute of Medical Microbiology and Hygiene, University of Regensburg, Regensburg, Germany; Institute of Clinical Microbiology and Hygiene, University Hospital Regensburg, Regensburg, Germany

**Keywords:** influenza, neuraminidase, pseudotype, ELLA, vaccine, monoclonal antibody, antisera, inhibition

## Abstract

To better understand how inhibition of the influenza neuraminidase (NA) protein contributes to protection against influenza, and to investigate its breadth and cross-neutralizing activity, we have produced lentiviral vectors pseudotyped with an avian H11 hemagglutinin (HA) and the NA (N1-N9) of all influenza A and (B/Victoria and B/Yamagata) influenza B subtypes. These NA viral pseudotypes (PV) possess stable NA activity and can be utilized as target antigens in *in vitro* assays to assess vaccine immunogenicity. Employing these NA PV, we have developed an enzyme-linked lectin assay (pELLA) for routine serology to measure neuraminidase inhibition (NI) titers of reference antisera, monoclonal antibodies, and post-vaccination sera with various influenza antigens. We have also shown that pELLA is more sensitive than the commercially available NA-Fluor™ in detecting NA inhibition in these samples. Our studies may lead to establishing the protective NA titer that contributes to NA-based immunity. This will aid in the design of superior, longer lasting, and more broadly protective vaccines that can be employed together with HA-targeted vaccines in a pre-pandemic approach.

## 1. Introduction

Influenza virions display two major surface glycoproteins, hemagglutinin (HA) and neuraminidase (NA) that play crucial roles in influenza infection and immunity. HA is involved in viral entry and NA in viral release (1–3). Currently available Influenza A (IAV) and Influenza B (IBV) vaccines mainly target HA, however, HA is subject to constant antigenic drift at distinct epitopes in its exposed globular head domain (4–6). As a result, seasonal influenza vaccines are reformulated annually to provide up to date protection.

Neuraminidase is a tetramer formed of separate subunits consisting of a head and stalk region, similar to the HA proteins (7). NA has known enzymatic activity which enables it to cleave sialic acid from cellular and viral glycoproteins expressed in infected cells (7). Sialic acids (SA) are typically found at the terminals of oligosaccharides on gangliosides and glycoproteins. SA are mainly linked to galactose residues by α-2,3 linkages, most commonly found in avian species, or α-2,6-linkages, mainly found in humans and other mammals (8). NA cleaves α-2,3 linked sialic acids more efficiently than α-2,6 sialic acids indicating that influenza NA is more specialized to avian infection (9,10). However, studies have shown that α-2,6 activity increases over time (11), which in combination with changes to HA may allow avian influenza strains to ‘species jump’ and potentially cause a pandemic in humans (12). Aside from its enzymatic activity, NA is vital in preventing the aggregation of viral progeny (13), enables penetration of human mucus by freeing virus from sialyated host mucins (2) and also facilitates viral budding (14). Neuraminidase (NA) is subject to antigenic changes, although at a lower rate than HA (15–17).

Neuraminidase has been a successful target for antiviral drugs and is currently being studied as an alternative or adjunct to HA as a viable vaccine candidate with the potential to be employed as part of a ‘universal’ or ‘cross subtype’ influenza vaccine (18–20). The ultimate goal of a universal vaccine is to protect against infection from novel influenza viruses bearing combinations of H1-H18 and N1-N11. However, this is a highly ambitious undertaking and consensus within the field is that influenza vaccines with approximately 75% protection against influenza A and B viruses that can protect for at least 12 months across a range of age groups and socio-economic backgrounds is considered an achievable interim aim (21). There are 144 possible combinations of NA and HA subtypes for non-bat IAV and 120 combinations have been observed in nature (22,23). Studies have shown that anti-NA antibodies raised following immunization with N1 (H1 subtype) can inhibit homologous and heterosubtypic influenza A viruses (e.g. H5N1, H3N2 & H7N9) (24) and are successful in controlling influenza infection *in vivo* (25). It is also of note that currently licensed inactivated influenza vaccines do contain NA and the quality and stability of NA in these preparations has not been fully investigated (26). It is known that the anti-NA seroconversion rate of individuals immunized with inactivated trivalent vaccine is variable (27–29) and this could potentially be increased by considering NA as an immunogen during vaccine formulation.

Antibodies directed to NA do not block viral entry and are therefore not classified as classically neutralizing antibodies (30). Traditionally, the inhibition of NA enzymatic activity has been measured using a MUNANA substrate-based assay (31,32), however this assay utilizes hazardous chemicals making it unsuitable for high throughput serological testing. Therefore, alternatives assays such as the Enzyme Linked-Lectin Assay (ELLA) (33–36) and fluorescence-linked MUNANA based assays (e.g. NA-star/NA Fluor™) have been developed to quantify NA enzymatic activity as a measure of antibody mediated inhibition of viral egress from infected cells (37). These methods make the study of NA activity more accessible for the design of the next generation of influenza vaccines. Additionally, pseudotype virus (PV) can also be used as a substitute to wild type virus in these assays. Neuraminidase pseudotyped viruses have already been successfully used in place of reassortant virus or Triton X-treated wild type virus in the ELLA assay for N1 and N2 subtypes (38,39) As NA has the potential to be included in a more broadly protective vaccine approach, a toolbox of assays capable of assessing NA inhibition will be required. To this end, we have produced an NA PV library encompassing IAV N1-N11 and IBV from the Victoria-like (B/Vic) and Yamagata-like (B/Yam) lineages for use in the pELLA assay and potentially the NA-Fluor™ assay. We demonstrate that these PV can be effectively employed in the pELLA to assess NA inhibition of reference anti-NA antisera, monocolonal antibodies (mAb), and anti-NA antibodies generated through vaccination, without the requirement for containment higher than BSL 2.

## 2. Materials and Methods

### 2.1 Production and transformation of plasmids

Neuraminidase genes from IAV subtypes N1-N9 and IBV, B/Vic, and B/Yam, were gene-optimized and adapted to human codon use via the GeneOptimizer algorithm (40). These NA genes were cloned into pEVAC (GeneArt, Germany) via restriction digestion. Plasmids were transformed via heat-shock in chemically induced competent *E. coli* DH5α cells (Invitrogen 18265-017). Plasmid DNA was extracted from transformed bacterial cultures via the Plasmid Mini Kit (Qiagen 12125). All plasmids were subsequently quantified using UV spectrophotometry (NanoDrop™ -Thermo Scientific).

### 2.2 Production of influenza H11-NA(X) pseudotypes (PV)

For production of H11-NA(X) PV, human embryonic kidney 293T/17 (HEK293T/17, ATCC: CRL-11268^a^) were maintained in complete medium, (Dulbecco’s Modified Essential Medium (DMEM) (PANBiotech P04-04510) with high glucose and GlutaMAX supplemented with 10% (v/v) heat-inactivated Fetal Bovine Serum (PANBiotech P30-8500), and 1% (v/v) Penicillin-Streptomycin (PenStrep) (Sigma P4333)) at 37°C and 5% CO_2_. Transfection was done as previously described (41). On the day prior to transfection, 4×10^5^ HEK293T/17 cells in complete DMEM were seeded per well of a 6-well plate. The next day, media was replaced, and cells were transfected using FuGENE® HD Transfection Reagent (Promega E2312) in Opti-MEM™ (Thermo Fisher Scientific 31985062) with the following plasmids: 10 ng NA encoding plasmid (pEVAC), 10 ng H11 encoding plasmid (A/red shoveler/Chile/C14653/2016 (H11) (pEVAC), 5 ng transmembrane serine protease 4 (TMPRSS4) encoding plasmid, 375 ng luciferase vector plasmid (pCSFLW), and 250 ng p8.91 gag-pol (Gag-Pol expression plasmid. For the H5 release assay, 10 ng A/Indonesia/5/2005(H5) (pI.18) was included in the plasmid DNA mixture replacing H11 (pEVAC), and TMPRSS4 was not utilized. All plasmid DNA were combined in 100 µL OptiMEM™, and FuGENE® HD (3 µL per µg plasmid DNA) was added dropwise followed by incubation for 15 minutes. The plasmid DNA-OptiMEM™ mixture was then added to the cells with constant swirling. Plates were incubated at 37°C, 5% CO_2_ for 48 hours. Supernatants were then collected, passed through a 0.45 μm filter, and stored at − 80°C.

### 2.3 H5 release assay

The ability of different NA to release H5 HA PV from producer cells was assessed by titration of the H5-NA(X) pseudotyped viruses produced as above in HEK293T/17 cells. Titration experiments were performed in Nunc F96 MicroWell white opaque polystyrene plates (Thermo Fisher Scientific 136101). Briefly, 50 µL of viral supernatant were serially diluted two-fold down columns of a 96-well plate in duplicate before adding 50 μL of 1×10^4^ HEK293T/17 cells to each well. No PV/cell only negative controls were included on each plate as an indirect cell viability measurement. Plates were then incubated at 37°C, 5% CO_2_ for 48 hours. Media was then removed and 25 µL Bright-Glo® luciferase assay substrate was added to each well. Titration plates were then read using the GloMax® Navigator (Promega) and the Promega GloMax® Luminescence Quick-Read protocol. Viral pseudotype titer was then determined as Relative Luminescence Units/mL (RLU/mL).

### 2.4 Reference antisera, monoclonal antibodies, and serum samples

Reference antisera to assess the inhibition sensitivity of representative IAV, anti-N1 A/California/7/2009 (NIBSC 10/218), anti-N1 A/turkey/Turkey/1/2005 (N1) (NIBSC 08/126), anti-N2 A/Victoria/361/2011 (NIBSC 14/144), and anti-N2 A/South Australia/34/2019 (NIBSC 19/320) antisera, and IBV, NA antiserum prepared from B/Malaysia/2506/2004 (NIBSC 05/252), and NA antiserum prepared from B/Florida/4/2006 (09/316), were obtained from the National Institute for Biological Standards and Control (NIBSC). Monoclonal antibodies against N1, 3A2, 1H5, 4E9, and 3H10 (42), as well as CD6 (43) and CR9114 (44) were also utilized in neuraminidase inhibition assays. Post-vaccination mouse sera were obtained from the University of Cambridge as part of an ongoing influenza vaccination study and were employed to determine responses of different NA antigens used for vaccination against corresponding H11_NA(X) pseudotypes.

### 2.5 Titration of NA PV via Enzyme Linked Lectin Assay (pELLA)

Clear Nunc Maxisorp™ flat-bottom 96-well plates (Thermo Fisher 44-2404-21) were coated overnight at 4°C with 100 µL per well of 25 µg/mL fetuin (Sigma F3385) in 1X KPL coating buffer (Sera Care 50-84-00) to assess H11_NA(X) PV neuraminidase activity via pELLA (**Figure 1**). The next day, plates were washed three times with wash buffer (WB) (0.5% (v/v) Tween-20 in PBS). Two hundred forty µL of H11_NA(X) PV was serially diluted two-fold from neat to 1:2048 with sample diluent (SD) (1% (w/v) Bovine Serum Albumin (BSA), 0.5% (v/v) Tween-20 in PBS) across a row of a 96-well mixing plate. Fifty µL of the PV dilutions were then transferred to two rows until column 10 of the fetuin-coated 96-well plate, with columns 11 and 12 containing only SD (no PV control). All wells were then added with 50 µL SD. Plates were incubated overnight at 37°C. The next day, plates were washed 6 times with WB. One hundred µL of conjugate (1 µg/mL lectin from *Arachis hypogaea* (peanut) peroxidase conjugate (Sigma L7759) in conjugate diluent (1% (w/v) BSA in PBS)) was added to all wells and plates were incubated at room temperature for 2 hours with shaking (225 rpm). Plates were then washed 3 times with WB before adding 100 µL 1-Step™ Ultra TMB-ELISA Substrate Solution (Thermo Fisher 34029) followed by incubation in the dark at room temperature with shaking (225 rpm) for 10 minutes. Reaction was stopped by addition of 100 µL 0.1 M H_2_SO_4_ per well. Optical density at 450 nm (OD_450_) was determined using the Tecan Sunrise™ microplate reader with Magellan™ data analysis software. Readings were normalized to 100% and 0% OD_450,_ and the dilution that resulted in 90% OD_450_ was selected as the PV dilution input for inhibition assays.

**Figure 1.**
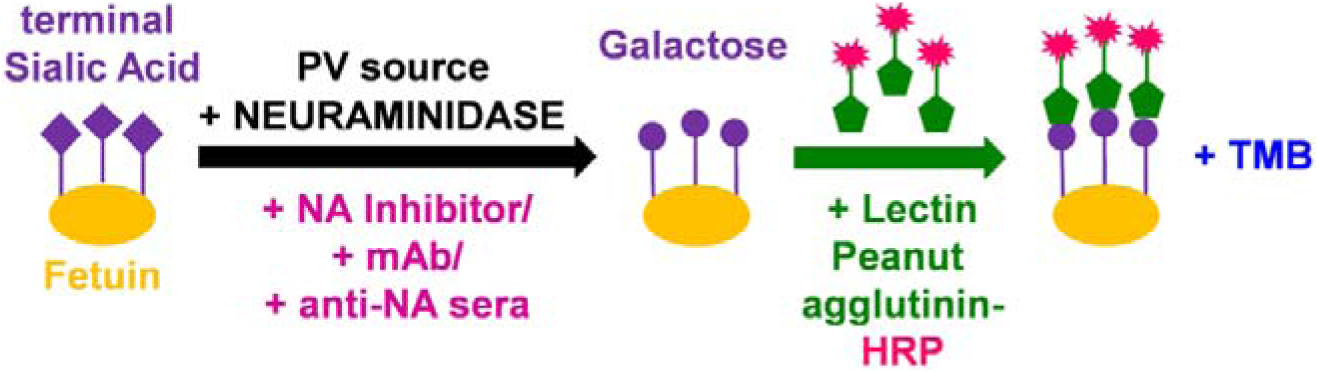
Cartoon illustrating main reactions in the pseudotype virus enzyme linked lectin assay (pELLA). The substrate fetuin is coated on each well of a 96-well plate. Neuraminidase, added via the NA PV we have generated, then cleaves the terminal sialic acid (SA) residues of the substrate fetuin giving rise to galactose residues. In the presence of substances that can impede NA such as mAbs and anti-NA antisera, the action of neuraminidase is inhibited and this can then be measured indirectly via pELLA. The terminal galactose residues that are present due to the action of neuraminidase are then specifically recognized by the substrate lectin peanut agglutinin conjugated to horseradish peroxidase (HRP). Addition of a peroxidase substrate such as TMB results in a detectable color change that can be measured at OD_450_.

### 2.6 Inhibition of NA PV by antisera and mAbs

The inhibition of neuraminidase activity by standard reference antisera, monoclonal antibodies (mAb), and serum samples from animal studies was evaluated via pELLA. Reference antisera were initially diluted 1:10 and then serially diluted five-fold; monoclonal antibody concentrations used were in the range of 32 µg/mL to 0.5 ng/mL. Serum samples from animal immunization studies were initially diluted 1:20 and then serially diluted two-fold in 50 µL SD. Dilutions were done in duplicate across two rows of a 96-well mixing plate. Fifty microliters of these dilutions were then transferred to fetuin-coated plates as described above, up to Column 10. Fifty µL of the H11_NA(X) PV that resulted in 90% OD_450_ as determined above (**Section 2.5**) was then transferred to all wells of the fetuin-coated plate except for Column 11 which served as the PV only control (0% inhibition) and column 12 contained SD only (100% inhibition). An additional 50 µL and 100 µL of SD was added to columns 11, and 12, respectively. All other steps were followed as per pELLA titration (**Section 2.5**). The IC_50_ was calculated as the inverse dilution of serum or antibody concentration that resulted in 50% inhibition of NA activity as determined via GraphPad Prism 9.0.

### 2.7 Flow Cytometry Binding Assay

HEK293T/17 cells were transfected with pEVAC encoding representative NA sequences of N1, N2, N3, N4, N8, and N9, as per PV production (**Section 2.2**). Forty-eight hours post-transfection, cells (50,000 cells/well) were transferred into V-bottom 96-well plates. Cells were then incubated with mouse sera (**Section 2.4**) (diluted 1:50 in PBS) for 30 minutes, washed with Fluorescence-activated cell sorting (FACS) buffer (PBS, 1% (v/v) FBS, 0.02% (v/v) Tween 20), and stained with goat anti-mouse IgG (H+L) Alexa Fluor 647 Secondary Antibody (Thermo Fisher A-21235) diluted at 20 µg/mL in FACS buffer, for 30 minutes in the dark. Cells were washed with FACS buffer and samples were run on the Attune NxT Flow Cytometer (Invitrogen) with a high throughput autosampler. Dead cells were excluded from the analysis by staining cells with 7-Aminoactinomycin D (7-AAD) and gating 7-AAD negative cells.

### 2.8 NA-Fluor™ Influenza Neuraminidase Assay

The NA activity of the H11-NA(X) PV was determined via the NA-Fluor™ Influenza Neuraminidase Assay (Life Technologies 4457091) kit. First, the linear range of fluorescence versus concentration of the Tecan Infinite 200Pro was determined by comparing relative fluorescence units (RFU) obtained from the NA activity assay of the PV to a standard curve of 4-Methylumbelliferone sodium salt 4-MU(SS) (Sigma M1508). NA activity is based on the production of 4-MU over time (60 minutes at 37 °C for the standard assay). The NA activity assay was then performed according to manufacturer’s protocol. Briefly, H11-NA(X) PV was diluted 2-fold from neat in NA-Fluor 2X assay buffer in a black 96-well plate and incubated for 1 hour at 37°C with the NA-Fluor substrate. After adding the stop solution, the plate was read at excitation and emission wavelengths of 355 nm and 460 nm, respectively, with an optimal gain of 55 using the Tecan Infinite 200Pro fluorescence plate reader. The RFU range for normalizing PV according to NA activity that is unique to every H11-NA(X) PV was then determined as well as the optimum H11-NA(X) PV dilution for the neuraminidase inhibition assay.

For the NA-Fluor™ Inhibition assay, the dilutions and concentrations for standard reference antisera, monoclonal antibodies (mAb), and serum samples from animal studies that were employed in the pELLA assay were likewise utilized (**Section 2.6**). The pre-determined amount of H11-NA(X) PV to test for inhibition of NA activity was then added. IC_50_ values were determined from dose-response data using sigmoidal curve-fitting generated by GraphPad Prism Software 9.0.

### 2.9 Statistical analyses

All statistical analyses were performed with GraphPad Prism 9 for Windows (GraphPad Software).

### 2.10 Bioinformatic analysis

NA sequences for both IAV and IBV were downloaded from the Influenza Virus Resource database (IVRD) (fludb.org). Phylogenetic tree was generated using the Cyber-Infrastructure for Phylogenetic RESearch (CIPRES) Gateway (45). The resulting tree file was then visualized using the Archaeopteryx tree viewer in the Influenza Resource Database (IRD) (46).

## 3. Results

### 3.1 NA PV production and functionality

Our NA PV library was constructed using the 4-plasmid transfection method as described previously (47) and encompasses NA IAV 1-11 and NA IBV from both B/Vic and B/Yam lineages (**Figure 2, Supplementary Table 1**). Similar to our HA PV library (47), p8.91 (48,49), a plasmid expressing the packaging genes (*gag-pol*) from a lentivirus (HIV) was used as the PV backbone. A pEVAC plasmid expressing the NA of the selected strain of IAV/IBV, pEVAC plasmid expressing hemagglutinin (HA) from A/red shoveler/Chile/C14653/2016 (H11), and a protease (transmembrane serine protease 4 (TMPRSS4)) to cleave HA were also included (**Figure 2A**). Employing this protocol, we have successfully produced representative H11-NA(X) pseudotyped virus for all IAV subtypes and both IBV lineages (**Figure 2B-C**).

**Figure 2.**
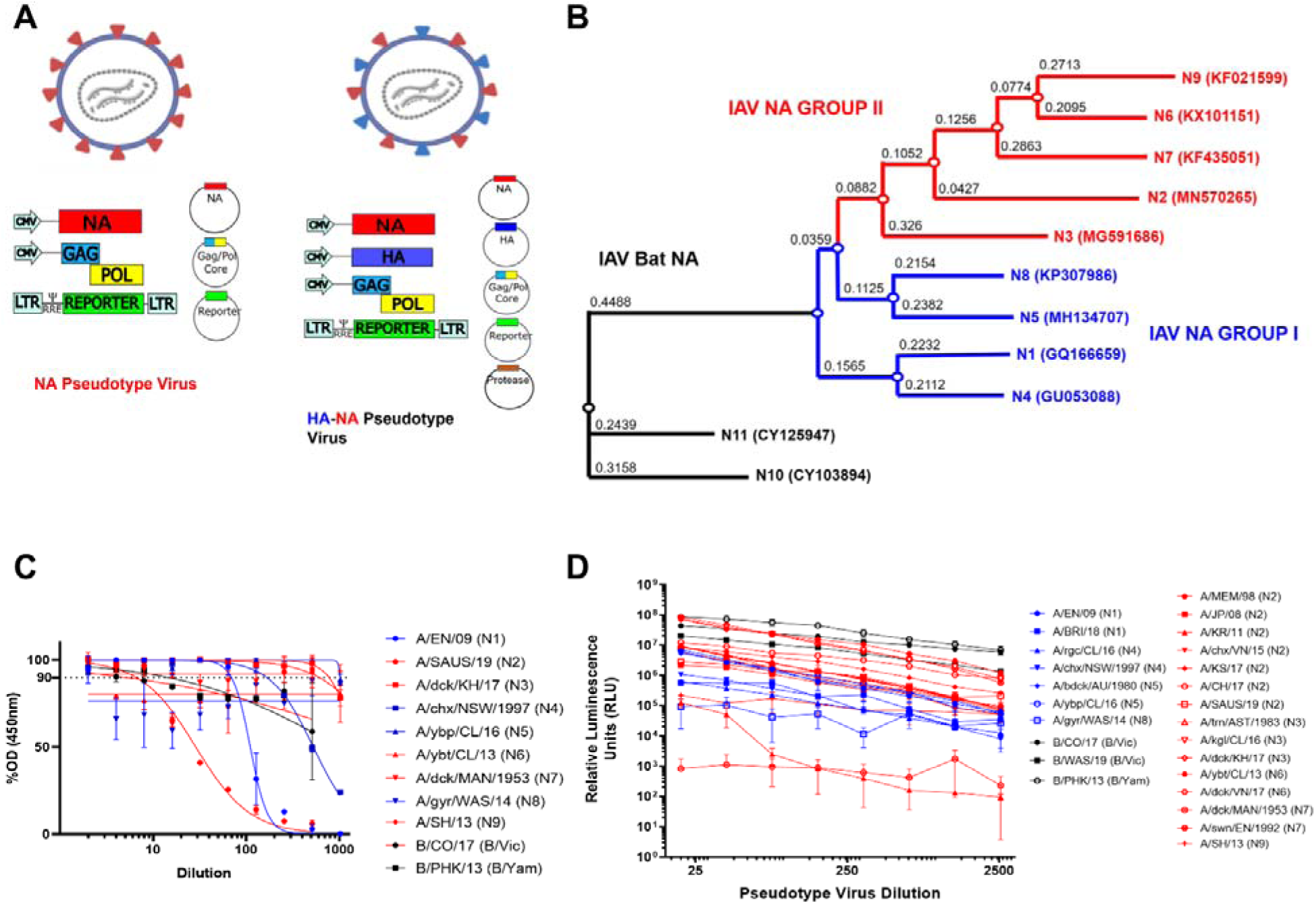
H11_NA(X) PV production and assessment of functionality. **(A)** Schematic representation of the production of influenza NA pseudotypes (PV) by quadruple plasmid transfection to produce pseudotypes expressing NA only or HA and NA on the PV surface. Protease is only required when HA is expressed, as per previous optimization (41). Images created using Biorender. **(B)** Phylogenetic tree of representative IAV NA from the PV library constructed. Influenza A Group I NA PV are shown in blue and IAV Group II PV in red. Accession numbers are reported with the subtype on the tree tips. Nodes are shown at the ends of branches which represent sequences or hypothetical sequences at various points in evolutionary history. Branch lengths indicate the extent of genetic change. The tree generated was constructed with PhyML on the Influenza Research Database (IRD) (46) and graphically elaborated with Archaeopteryx.js **(C)** Titration of Influenza A NA (1-9) and Influenza B NA (B/Victoria-like and B/Yamagata-like lineages) PV. Titration was carried out via pELLA. Readout is NA enzymatic activity expressed as %OD (450nm) of highest dilution tested. **(D)** Titration of Influenza A NA (1-9) and Influenza B NA (B/Victoria-like and B/Yamagata-like lineages) PV expressing H5 (A/Indonesia/05/2005) hemagglutinin. Readout is expressed in relative luminescence units (RLU). For **(C)** and **(D)**, each point represents the mean and standard deviation of two replicates per dilution (n=2). Additionally, Influenza A (IAV) Group I NA PV are shown in blue (N1, N4, N5, and N8), IAV Group II PV in red (N2, N3, N6, and N9), and influenza B NA PV (B/Victoria-like and B/Yamagata-like lineages) are shown in black.

We then assessed the neuraminidase enzymatic activity of the pseudotypes we produced via our optimized PV Enzyme Linked Lectin Assay (pELLA) protocol (**Figure 1**). ELLA pseudotype titrations were performed to determine the dilutions of PV required for pELLA inhibition assays. The PV was diluted two-fold from a starting dilution of 1:2 to 1:1024 and optical density at 450 nm was read (**Figure 2C**). All values were normalized against the highest PV value obtained (100% OD_450_) and the sample diluent only control (0% OD_450_). Traditionally, 90% activity is optimal for use in the inhibition assay (33,39), however, we also set the additional criteria of the OD_450_ to be a minimum of 2.0 for use in pELLA inhibition. All human and avian PV produced demonstrated NA enzymatic activity and met the additional criteria we set for use in pELLA inhibition (**Figure 2C**) apart from bat N10 and N11. These bat NA have been reported to not demonstrate any “classic” neuraminidase activity in these assays (50–54). We then utilized this PV library to test for anti-NA activity in post-vaccination mouse sera.

To assess the ability of our NA to release HA, we produced PV with IAV H5 and IAV NA (N1-N9) and IBV NA (B/Vic and B/Yam) in place of exogenous NA and titrated the PV via a luciferase reporter assay (41). N10 and N11 have previously been shown to not be required for HA release (41,55) and were therefore not tested herein. All the NA PV tested were capable of releasing H5 as evidenced by the titers observed (**Figure 2D**). H5 HA was selected as it is a highly pathogenic IAV subtype harboring a functional polybasic cleavage site that does not require an exogenous protease for HA release (56,57). Even though H5 has not been observed *in vivo* in combination with all NA subtypes utilized here, all PV produced RLU titers of 10^5^ and higher except for A/duck/MAN/1953 (N7) which data here suggests does not release H5 as efficiently (**Figure 2D**). Interestingly, one of the N2 PV constructed (A/Korea/KUMC_GR570/2011) only achieved a production titer of 10^5^, reasons for these discrepancies are not clear.

### 3.2 *In vitro* inhibition of NA pseudotypes by antisera and monoclonal antibodies

The inhibition susceptibility of representative NA PV generated to available NA subtype specific reference antisera was tested. We have shown here dose response curves for PV which have previously infected humans: N1 (A/England/195/2009), N2 (A/South Australia/34/2019), B/Vic lineage NA (B/Colorado/6/2017) and B/Yam lineage NA (B/Phuket/3073/2013) **(Figure 3A**). All reference antisera were able to inhibit the PV they were tested against (**Figure 3A-B**) including both anti-N1 2009 pandemic antisera, A/California/7/09(H1N1), and antisera A/turkey/Turkey/05, originally from A/turkey/Turkey/1/05 (H5N1), with IC_50_ dilution values of ∼25,000 and ∼1,300 respectively. Both N2 PV were inhibited by anti-N2 antisera, with IC_50_ dilution values of ∼32000 and ∼1,300. Similarly, IBV lineage PV were inhibited by anti-sera tested with IC_50_ dilution values of ∼6300 for B/Vic-like ∼ 39,000 for B/Yam-like antisera.

**Figure 3.**
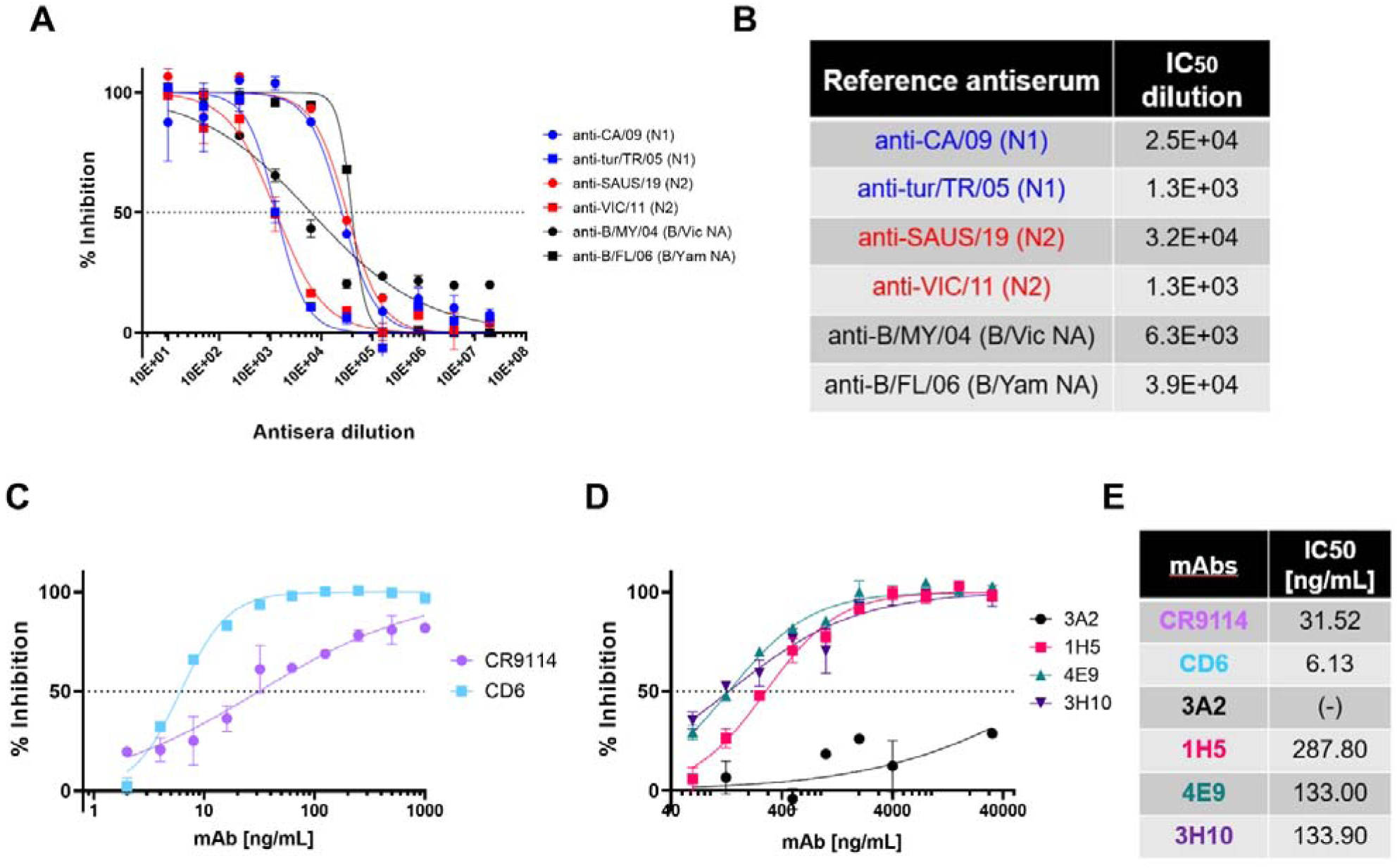
*In vitro* inhibition of NA pseudotypes by antisera and monoclonal antibodies. **(A)** *In vitro* inhibition of representative influenza A and influenza B NA pseudotype virus (PV) which have previously caused human infection: N1 (A/England/195/2009), N2 (A/South Australia/34/2019), B/Victoria-like lineage NA (B/Colorado/6/2017), and B/Yamagata-like lineage NA (B/Phuket/3073/2013) by reference antisera obtained from NIBSC. Reference antisera were serially diluted five-fold from a starting dilution of 1:10 and PV diluted to 90% OD_450_ as determined previously from titration (**Figure 2C**). Inhibition of NA activity was determined via pELLA. Each points represents the mean and standard deviation of two replicates per dilution (n=2). **(B)** Half-maximal inhibitory dilution of reference antisera as calculated from dose response curves (**A**). **(C)** *In vitro* inhibition of representative N1 (A/England/195/2009) PV by monoclonal antibodies CR9114 and CD6. mAbs were diluted two-fold from 1000 ng/mL to 2ng/mL. **(D)** *In vitro* inhibition of representative N1 (A/England/195/2009) PV by N1-directed mAbs: 3A2, 1H5, 7E9, and 3H10. mAbs were diluted 2-fold from 32 µg/mL to 32 ng/mL. For **(C)** and **(D)**, inhibition was determined via pELLA and each points represents the mean and standard deviation of two replicates per dilution (n=2). **(E)** IC_50_ concentration values as determined from **(C)** and **(D)** using non-linear regression.

IAV H1N1 has caused pandemic influenza in humans, most recently in 2009 (58,59). Taking this into account, we tested representative N1 PV (A/England/195/2009) inhibition sensitivity to a variety of monoclonal antibodies (mAb) in pELLA inhibition. As the PV created herein express H11, we tested the ability of HA stem-directed mAb CR9114 (44), which was previously shown to neutralize IAV and IBV HA PV (41), to inhibit NA enzymatic activity in a traditional ELLA (60). CR9114 inhibited the N1 PV tested (**Figure 3C**) with half maximal inhibitory concentration IC_50_ of 31.52 ng/mL (**Figure 3E**) as determined by non-linear regression. As CR9114 inhibits the action of NA via steric hindrance, we also investigated the inhibitory action of anti-N1 directed antibodies. We tested CD6, a mAb raised against H1N1 2009 pandemic strain of IAV that binds to an N1 conserved epitope which spans the lateral face of the NA dimer (61). We show here that our N1 PV (A/England/195/2009) was inhibited by CD6 (**Figure 3C**), with IC_50_ determined to be 6.13 ng/mL, more potent than CR9114. We also explored inhibition of our 2009 pandemic N1 PV with anti-N1 mAbs generated against A/Brisbane/59/2007 (42). Monoclonal antibodies 1H5, 4E9, and 3H10 (**Figure 3D**) had IC_50_ values of 287.80 ng/mL, 133 ng/mL, and 133.90 ng/mL, respectively (**Figure 3E**). However, mAb 3A2 did not achieve 50% inhibition (**Figure 3D-E**). Reasons for this and the high concentrations of mAb required, compared to CD6, may be explained by the PV utilized herein being a 2009 pandemic H1N1 strain. The mAb 3A2 was raised against a pre-pandemic strain, suggesting that the 3A2 specific epitope may have been affected by the antigenic shift seen in 2009.

### 3.3 Inhibition of neuraminidase activity by post-NA vaccination mouse sera

We obtained mouse sera from pre-clinical influenza vaccination studies in mice at the University of Cambridge and tested the ability of these mouse sera to bind and neutralize H11-NA(X) PV from our library representing all IAV subtypes and IBV lineages. Naïve mice were vaccinated with NA (IAV N1-N9 and IBV B/Vic and B/Yam) to elicit an immune response against influenza neuraminidase (**Figure 4**). Mouse terminal bleeds were obtained and were assessed via FACS for the capacity to bind to HEK293T/17 cells transfected with pEVAC encoding the homologous IAV NA subtype. As expected, high binding activity of post-vaccination sera to HEK293T/17 cells expressing the homologous NA was observed with log MFI values shown to be ≥4 (red in the heat map) (**Figure 4A**). Interestingly, sera from mice vaccinated with N2 and N4, bound to all NA expressed in HEK cells with log MFI values ranging from ∼3-4 (**Figure 4A**). Sera from mice vaccinated with N5, N6, and N7, displayed little to no binding to all the NA tested, while sera from mice vaccinated with N8 and N9 only had strong binding activity against homologous NA expressed in HEK cells (**Figure 4A**). Surprisingly, all sera from vaccinated mice showed modest cross-binding activity with N9 in HEK cells regardless of NA subtype they were vaccinated with.

**Figure 4.**
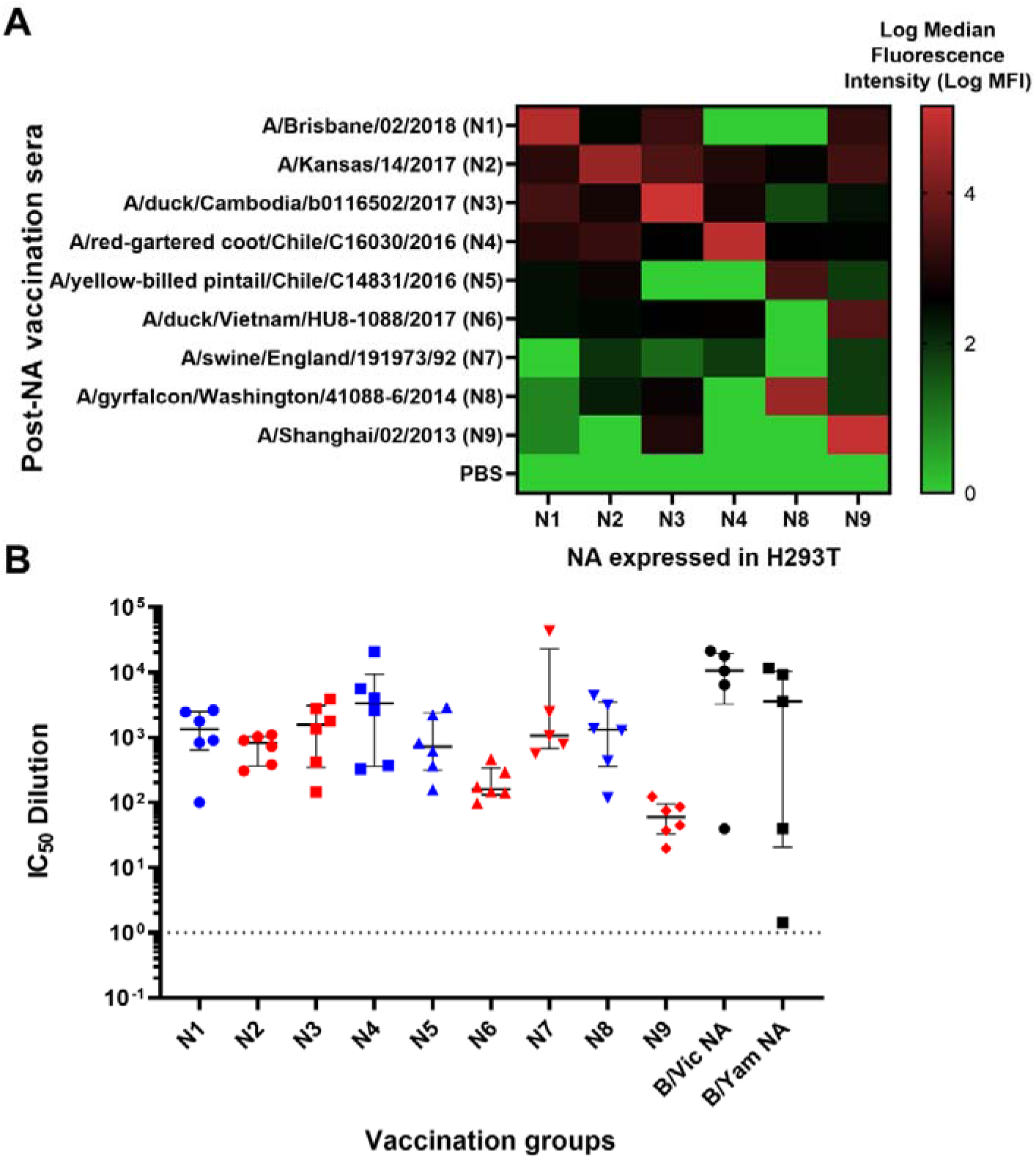
Binding and anti-NA activity of post NA vaccination sera. **(A)** Binding of representative post-NA vaccination sera to HEK293T/17 transfected with pEVAC encoding homologous neuraminidase sequences of N1, N2, N3, N4, N8, and N9, was determined via FACS and reported as Median Fluorescence Intensity (MFI) in a heatmap. Readings were done in duplicate (n=2). **(B)** *In vitro* inhibition of representative IAV (homologous subtype) and IBV (homologous lineage) pseudotypes by mouse sera vaccinated with Influenza A HA from A/Brisbane/2/18 (N1), A/Kansas/14/17 (N2), A/duck/Cambodia/b0116502/17 (N3), A/chicken/NSW/1688/1997 (N4), A/yellow-billed pintail/Chile/C14831/16 (N5), A/yellow-billed teal/Chile/8/13 (N6), A/swine/England/191973/1992 (N7), A/gyrfalcon/Washington/41088-6/14 (N8), and A/Shanghai/2/13 (N9), and Influenza B HA from B/Colorado/6/17 (B/Vic) and B/Phuket/3073/13 (B/Yam). Inhibition was determined via pELLA and reported as IC_50_ dilution values (IC_50_ is half maximal inhibitory serum dilution). For mice vaccinated with N1-N6, N8-N9, n=6, for N7 and both B lineages, n=5. Group I NA are indicated in blue and Group II NA in red. Plot shows the median and interquartile range of all samples tested.

We then performed pELLA inhibition employing these post-vaccination mouse sera against NA PV from our library to assess NA vaccine immunogenicity. All post-vaccination sera neutralized the homologous NA subtype (IAV)/lineage (IBV) pseudotype they were tested against with IC_50_ dilution values ranging from ∼100 to ∼10,000 (**Figure 4B**). These results indicate that post-vaccination immune responses can be effectively evaluated using our optimized pELLA.

### 3.4 Comparison of pELLA with NA-Fluor™ to evaluate NA activity and inhibition

We then compared the results of NA activity and inhibition as determined by pELLA to that obtained via the NA-Fluor™ assay, a commercially available fluorescence linked-MUNANA based assay routinely used to monitor neuraminidase inhibitor (NI) drug sensitivity.

First, we determined the linear range of fluorescence versus concentration of fluorescent 4-Methylumbelliferone (4-MU) of the instrument, the Tecan Infinite 200Pro. 4-MU(SS) is the end product when the substrate, 20-(4-methylumbelliferyl)-a-D-N-acetylneuraminic acid (MUNANA), is cleaved by NA (31,62–64). We then selected an RFU value within the linear range of fluorescence detection of the instrument for normalizing PV according to NA activity (**Supplementary Figure 1**). From these findings (**Supplementary Figure 1)**, we then chose 10,000 RFU (shown via broken line) as the fluorescence signal output for NA activity normalization (**Figure 5A**). For each PV, we then selected the dilution factor that yielded 10,000 RFU as identified in the 4-MU(SS) standard curves. Our results show that we have successfully produced H11-NA(X) PV with neuraminidase activity that is within the linear dynamic range of detection of the NA-Fluor™ assay that we utilized (**Figure 5A**). These results corroborate the NA activity of our PV library as determined via pELLA (**Figure 2C**).

**Figure 5.**
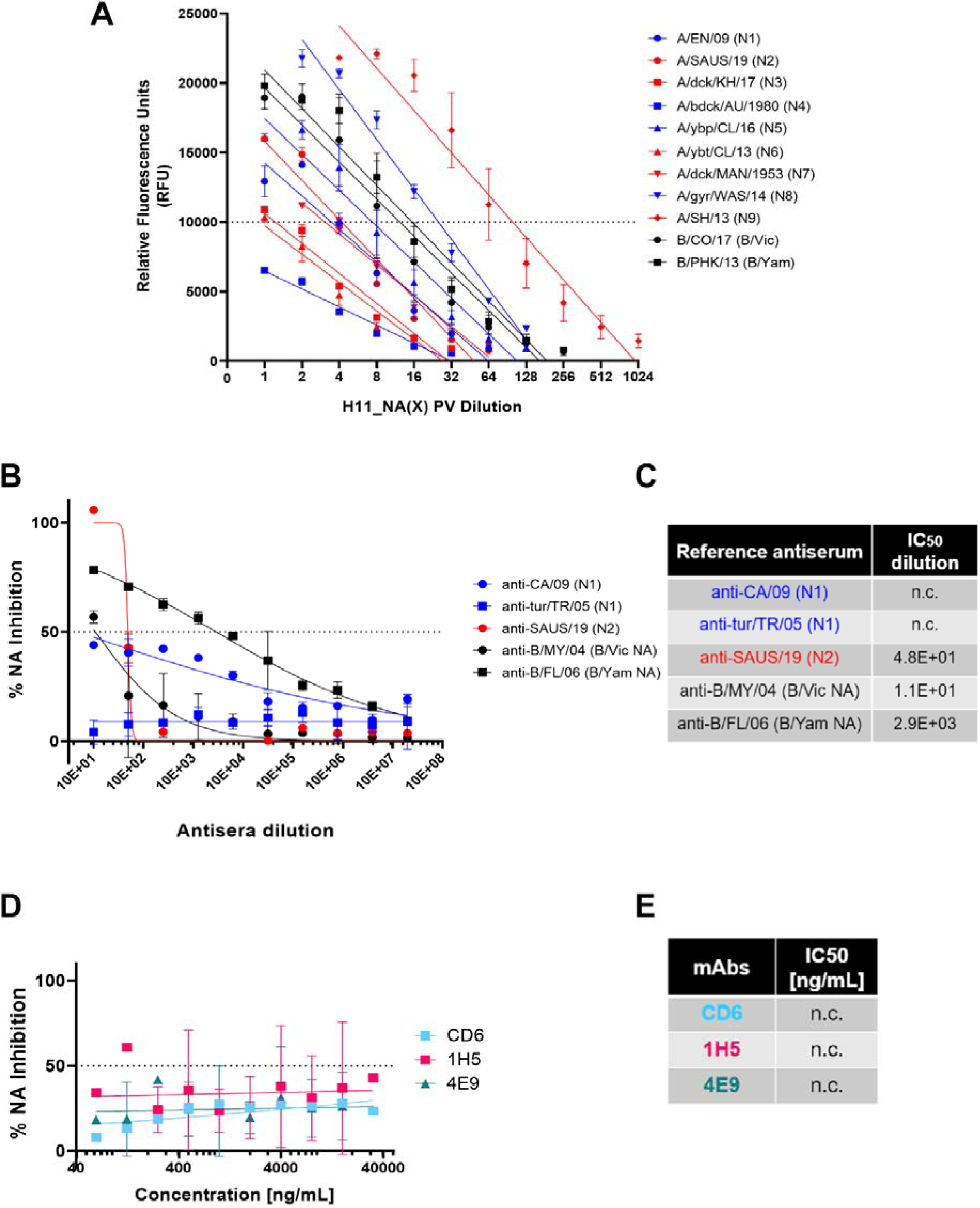
Determination of NA activity of H11_NA pseudotypes and NA inhibition activity of reference antisera and mAbs via NA Fluor™. **(A)** Titration of representative IAV (N1-N9) and IBV (B/Victoria-like and B/Yamagata-like lineages) PV. Titers are reported in Relative Fluorescence Units (RFU) (n=2). Dotted line at 10,000 RFU indicates value used for NA activity normalization (**Supplementary Figure 1**). **(B**,**C)** Inhibition of A/England/195/09 (N1) (blue), A/South Australia/34/19 (N2) (red), IBV (black) B/Colorado/6/17 (B/Vic NA) and B/Phuket/3073/13 (B/Yam NA) H11_NA PV by reference antisera. **(B)** Reference antisera were serially diluted five-fold from a starting dilution of 1:10, similar to **Figure 3A**. NA PV at a dilution that would give 10,000 RFU as determined in **(A)** was then added to each well. **(C)** IC_50_ dilutions for reference antisera against homologous subtype/lineage PV are shown. For **A-C**, Group I NA PV are shown in blue, Group II NA PV in red, and IBV (both lineages) in black. **(D-E)** Inhibition of A/England/195/09 (N1) PV by N1-specific monoclonal antibodies. **(D)** Monoclonal antibodies were serially diluted two-fold from a starting concentration of 32 µg/mL to 0.0625 ng/mL, and IC_50_ values are summarized in **(E)**. For plots **(A), (B)**, and **(D)**, each point represents the mean and standard deviation of two replicates per dilution. For **(C)** and **(E)**, “n.c” indicates values not computed by GraphPad Prism.

We then attempted to demonstrate NA inhibition of our representative NA PV by the same reference antisera and monoclonal antibodies we tested previously in the pELLA (**Figure 3**). Despite employing the same reference antisera dilutions (**Figure 5B-C**) and mAb concentrations (**Figure 5D-E**), we did not observe the same inhibition activity against the PV tested in the NA-Fluor™ assay. For the reference antisera, only the anti-B/FL/06 (B/Yam NA) showed appreciable neutralizing activity against the representative B/Yam NA PV (**Figure 5B**), however, the IC_50_ dilution value determined here (**Figure 5C**) was a log lower from that observed in the pELLA (**Figure 5B**). The rest of the reference antisera tested did not seem to strongly inhibit their PV counterparts (**Figure 5B-C**). Similar to the reference antisera, the mAbs specific against N1 that were tested did not inhibit A/England/195/2009 (N1) PV (**Figure 5D-E**), with the data generated insufficient to calculate IC_50_ values (n.c).

We then compared NA inhibition via pELLA and NA-Fluor™ by post-NA vaccination mouse sera (**Figure 6**). We employed the same post-vaccination serum samples and tested using the same PV, yielding contrasting results. NA neutralization of the N1 and N2 PV employed was successfully demonstrated using the pELLA for all tests subjects while there was no inhibition observed using the NA-Fluor™ (**Figure 6**).

**Figure 6.**
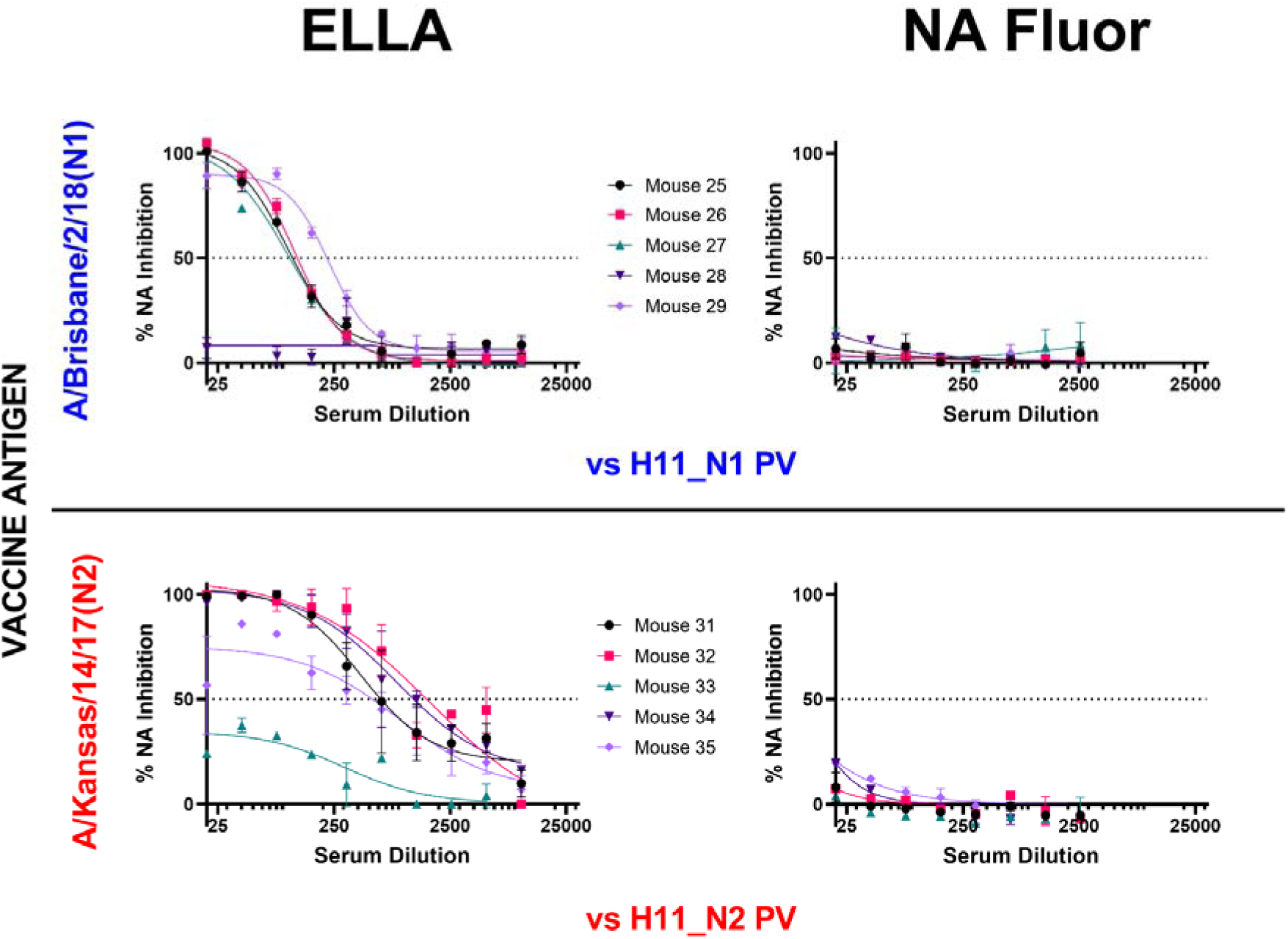
Comparison of inhibition of H11_NA PV by N1 and N2 post-vaccination mouse sera using enzyme linked lectin assay (pELLA) and NA Fluor™. Sera from mice vaccinated with A/Brisbane/2/18 (N1) (blue) (n=6) and A/Kansas/14/17 (N2) (red) (n=6) were diluted two-fold starting from 1:20 for both assays. They were then tested for their ability to inhibit homologous NA subtype PV. Percent NA inhibition is then shown as a function of serum dilution. For all plots, each point represents the mean and standard deviation of two replicates per dilution.

## 4. Discussion

Development of a Universal Influenza Vaccine relies heavily on the determination of specific immune responses and other correlates of protection that prevent illness due to influenza infection (26,65,66). Seasonal influenza vaccination targets the influenza HA head which has been shown in the past to be the most immunodominant influenza antigen, with NA, the second most abundant influenza surface glycoprotein, being overlooked until recently (66). Neuraminidase performs multiple functions in the influenza infection cycle which gives rise to the many possible avenues where antibodies against NA can be exploited to provide protection against influenza. Similarly to HA, antibodies specific to the conserved regions of NA have shown excellent breadth and are able to inhibit divergent influenza viruses (60,67).

Although limited, knowledge about immunity based on inhibition of NA activity has been building up over the years. Several clinical studies have successfully shown the importance of the presence of strong anti-NA inhibition titers (either vaccination-induced or pre-existing through natural infection) in decreasing the frequency of influenza infection and illness (68–70). These NA inhibition titers (NAI) were found to be independent of hemagglutination inhibition (HAI) titers, both of which can be used in conjunction to determine influenza sero-protection (68). These findings warrant further studies on anti-NA immunity by applying similar methods that allowed us to study anti-HA responses, solidifying NA’s important role in influenza prevention and control.

The function of NA and interactions *in vivo* with HA are increasingly being explored especially in relation to the design and efficacy of novel vaccine or drug candidates. The NA PV library we have constructed can be utilized to this end in a similar fashion to the HA PV library we previously described (41). Our comprehensive library has representative strains of IAV N1-11 and IBV B/Vic NA and B/Yam NA (**Supplementary Table 1)** and the methods used to produce these PV can easily be employed to include additional NA strains as required. The PV library could be applied to research for human disease with emphasis on the zoonotic potential of strains such as H10N3 reported recently in China (71) that may have pandemic potential. We produced NA PV pseudotyped with an HA (in this case, H11) although it was possible to produce NA PV on its own (**Figure 2A**), as we found that when testing inhibition capacity of mouse sera, an HA plasmid was required to maintain stability of the PV as observed previously (72) and reduced background for use in pELLA (34,38). Additionally, this arrangement mirrors the influenza wild type virus surface that contains both HA and NA, potentially providing a more accurate model for interactions between these surface glycoproteins that can then be probed at lower containment.

All IAV PV (N1-N9) and both lineages of IBV (Victoria-like and Yamagata-like lineages) showed NA enzymatic activity in our optimized pELLA (**Figure 2C**) suggesting that the PV produced herein have functional NA sialidase activity. We have also shown that most of the NA we tested were able to release the H5-NA(X) PV (**Figure 2D**). However, we have reported an unusual case, where A/duck/MAN/1953 (N7) that had a lower H5 release capacity compared to the other NA tested (**Figure 2D**) has demonstrated strong neuraminidase activity (**Figure 2C**). This may be because this particular H5 from A/Indonesia/5/2005 has not been shown to combine in nature with all the NA subtypes tested herein. Furthermore, our data implies that the role NA plays in HA release (**Figure 2D**) and NA enzymatic activity, as measured via pELLA (**Figure 2C**), may be more complex than a direct correlation between enzymatic activity and HA release capacity and therefore warrants further investigation (73).

There is a limited number of subtype specific anti-NA antisera available for testing as there are only very few NA subtypes that have infected humans in the past, nonetheless, we have shown that the antisera we obtained could inhibit homologous subtype NA activity in the pELLA (**Figure 3A-B**). For mAbs, we utilized CR9114, a broadly-neutralizing anti-HA stem antibody that binds in a structurally similar manner to the HA stem region across HA subtypes 1-16, blocking the HA pH-induced conformational changes associated with membrane fusion (44). HA-stem specific antibodies like CR9114 inhibit NA activity via steric hindrance (60) and detection of this activity this may be critical for the advancement of universal vaccines which utilize similar targets (24,74–76). The data we presented (**Figure 3C**) shows CR9114 inhibiting the NA enzymatic activity of H11-N1 PV, affirming that the pELLA can also be employed to detect this activity similar to traditional ELLA assays (60). Additionally, we demonstrated that the inhibiting activity of mAbs which directly target epitopes of NA can also be assessed via this assay (**Figure 3C-D**). Although we only had access to a limited selection of anti-NA N1 mAbs we have demonstrated that our H1N1 pandemic strain (A/England/195/2009) was preferentially inhibited by CD6, a mAb raised against H1N1 2009 pandemic strain (A/California/7/2009) compared to mAbs raised against the seasonal influenza strain (A/Brisbane/59/2007) (**Figure 3C-D**). These findings concur with previous experiments conducted with traditional pELLA assays by Wan et al (43).

A broadly protective vaccine directed at NA epitopes is becoming an increasingly plausible option. Previous studies have established that NAI antibody activity is an independent correlate of protection in humans (69,70). The breadth of immunity due to NA has not been fully elucidated, however a pertinent example of NA-based immunity can be gleaned from the 1968 H3N2 pandemic. The H3N2 pandemic virus of 1968 replaced the previous pandemic H2N2 subtype, as the H2 and PB1 gene segments were replaced by reassortment with an avian-like H3 HA and PB1, with its NA remaining the same (77). This pandemic H3N2’s sporadic nature and milder impact in terms of morbidity and mortality compared to its ancestors in different regions of the world is hypothesized to be mediated by N2 immunity from the previous pandemic in all age groups and to the HA antigen in the elderly (17,78,79). Similarly, NA-mediated immunity between H1N1 and H5N1 viruses has also been observed and this is due to conserved N1 epitopes between these viruses (24,25,61,67,80). Furthermore, mAbs isolated from H3N2 infected donors which bind directly to the NA active site have demonstrated exceptional breadth against NA from IAV and IBV and are broadly protective *in vivo* (81). This indicates a molecular basis for protection, and that these conserved protective epitopes can be ideal targets for a broadly protective vaccine.

We further studied NA-based protection by immunizing groups of mice with different NA subtypes (N1-N9) and looking for any cross-reactivity that may be present within subtypes. Results were interesting, as antibodies in post-vaccination mouse sera were able to bind to heterologous NA subtypes (**Figure 4A**) alluding to the probable presence of conserved epitopes among these subtypes, especially in mice vaccinated with N1 and N4, which are the closest relatives in the NA Group I phylogenetic tree (**Figure 2B**). Surprisingly, mouse post-vaccination sera bound to H293T cells expressing N2 and N9 (**Figure 4A**). Employing pELLA, we were able to successfully quantify NA inhibition in post-vaccination mouse sera, an indirect measure of antibodies against NA (**Figure 4B**). These findings are the first steps in identifying conserved regions or epitopes hidden in NA, followed by elucidating the mechanisms by which these NA antibodies contribute to protection and how these NAI antibody titers may be considered protective.

We then compared the results we obtained from pELLA with the NA-Fluor™ Influenza Neuraminidase Assay Kit. The NA-Fluor™ is a fluorescence-based assay, which quantifies the fluorogenic end product 4-methylumbelliferone released from the non-fluorescent substrate 2’-(4-methylumbelliferyl)-α-D-N-acetylneuraminic acid (MUNANA) by the enzymatic activity of neuraminidase. The amount of fluorescence directly relates to the amount of enzyme activity. This assay is widely accepted for monitoring of the effect of NA Inhibitors on NA activity or monitoring NI sensitivity in cell-based virus growth or inhibition assays (26). Employing the NA-Fluor™, we have shown that we produced H11-NA(X) PV with neuraminidase activity (**Figure 5A**) that can be used in NA-Fluor™ inhibition assays.

It is known that NA activity can be inhibited by antibodies binding directly to epitopes within the enzyme active site or through steric hindrance when antibodies bind proximal to the active site. However, NI through steric hindrance can only be observed when larger substrates, such as fetuin, are used, as in ELLA (81,83–85), and this may not be the case when smaller molecules are used as substrate as in the NA-Fluor™ and as such, antibodies that do not bind directly to the active site might not inhibit NA activity. We were also unable to show any NA inhibition activity in post-NA vaccination mouse sera (**Figure 6**), even if they all strongly inhibited NA as seen via the pELLA (**Figure 4B**). These results are somewhat disappointing as we could not find any correlation between the two methods and will require further investigation in the future.

Nonetheless, for the first time to our knowledge, we have demonstrated the utility of our optimized PV enzyme-linked lectin assay (pELLA) employing neuraminidase pseudotyped viruses to evaluate serologic responses to vaccination with IAV NA1-NA9 and IBV B/Victoria-like and B/Yamagata-like lineage NA. The pELLA was also useful in determining anti-NA directed monoclonal antibody and anti-NA reference antisera activity. The assay as it is shown here is more accessible to laboratories without high containment facilities, and low-resource environments than traditional MUNANA assays. The pELLA was also able to detect NA inhibition by a variety of samples more effectively than the commercially available NA-Fluor™.

Neuraminidase is an important protein which is now recognized to have multiple functions beyond its key role in viral budding during influenza infection (14) including: preventing aggregation of viral progeny (13), neutralizing protective effects of human mucus (86), and is capable of replacing functions of HA involved in viral entry (87–89). In the case of bat IAV, HA uses the major histocompatibility complex (MHC) for entry (55) and therefore it follows that N10 and N11 have not demonstrated sialidase activity (50–53). Recent data have shown that N11 is capable of downregulating cell surface expression of MHC-II molecules, the exact mechanism for this has not yet been elucidated (54). Currently there are several licensed drugs and small molecules which target NA (90), such as neuraminidase inhibitors (NAI) (91), strongly suggesting that a vaccine capable of eliciting anti-NA responses would be beneficial to preventing disease from influenza. Additionally, observations from natural immunity, including the broadly protective mAbs recently described (81) and a growing number of recombinant NA-based vaccines demonstrating protection in animal models (24,83,92,93) strengthen the need for further studies of NA as a viable vaccine target. In order to properly assess the immunity provided by anti-NA vaccines, a toolbox of assays will be required to predict *in vivo* protective efficacy and establish correlates of protection. We propose that the pELLA system described herein can form an important part of this new generation of *in vitro* vaccine assessment options. The flexibility of the PV production process ensures that immunity to a range of subtypes and strains can be tested against heterologous NA in a format that displays surface glycoproteins in their native conformation.

## Author Contributions

Conceived and designed experiments – KdC, JMD, GC, MF, NT; Performed experiments – KdC, JMD, MF, GC; Plasmid construction: SV, BA, RW; Analayzed the data – KdC, JMD, MF, NT; Reagent provision – RK, NT, JH; Wrote the paper – KdC, JMD, Revised the paper – GC, JH, RW, NT

## Acknowledgments

We would like to thank Jerry Weir of the US Food and Drug Administration (10903 New Hampshire Avenue, Silver Spring, MD 20993, USA) for the anti-N1 monoclonal antibodies we used in this study.

## 6. Supplementary Material

**Supplementary Figure 1.**
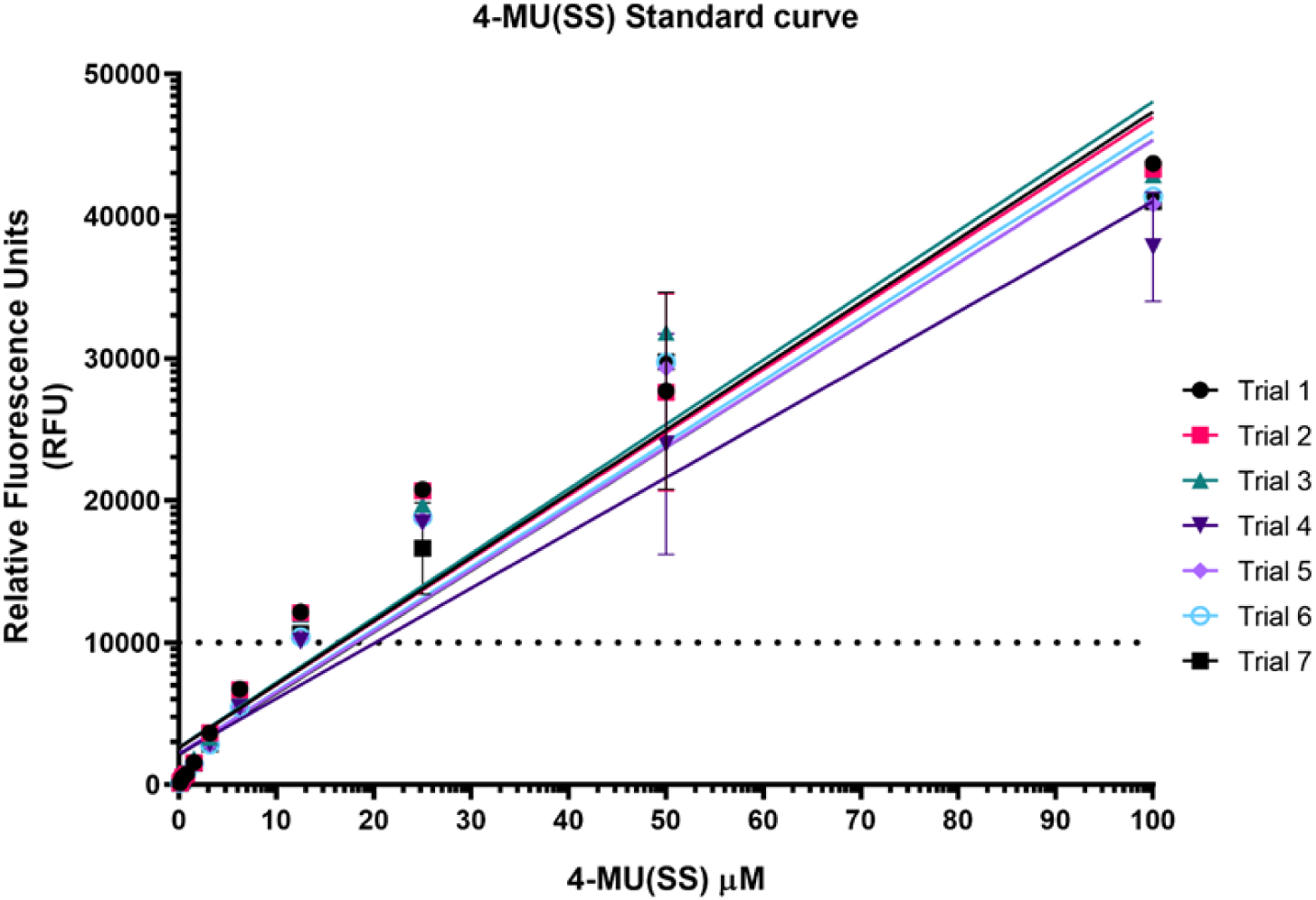
Standard Curve of 4-Methylumbelliferone sodium salt (4-MU(SS)). A standard curve was generated with different concentrations of 4-MU(SS) corresponding to a range of RFU values as read via the Tecan Infinite 200Pro fluorescence plate reader at excitation and emission wavelengths of 350 nm and 440 nm respectively (n=7). The Neuraminidase (NA) activity/RFU range for the NA inhibition assay was determined using the linear range of the 4-MU(SS) standard curve. We have arbitrarily chosen ∼12 μM 4-MU(SS) corresponding to ∼10,000 RFU (broken line) to normalize the NA activities of each PV for use in the NA-Fluor™ NA inhibition assay.

**Supplementary Table 1.**
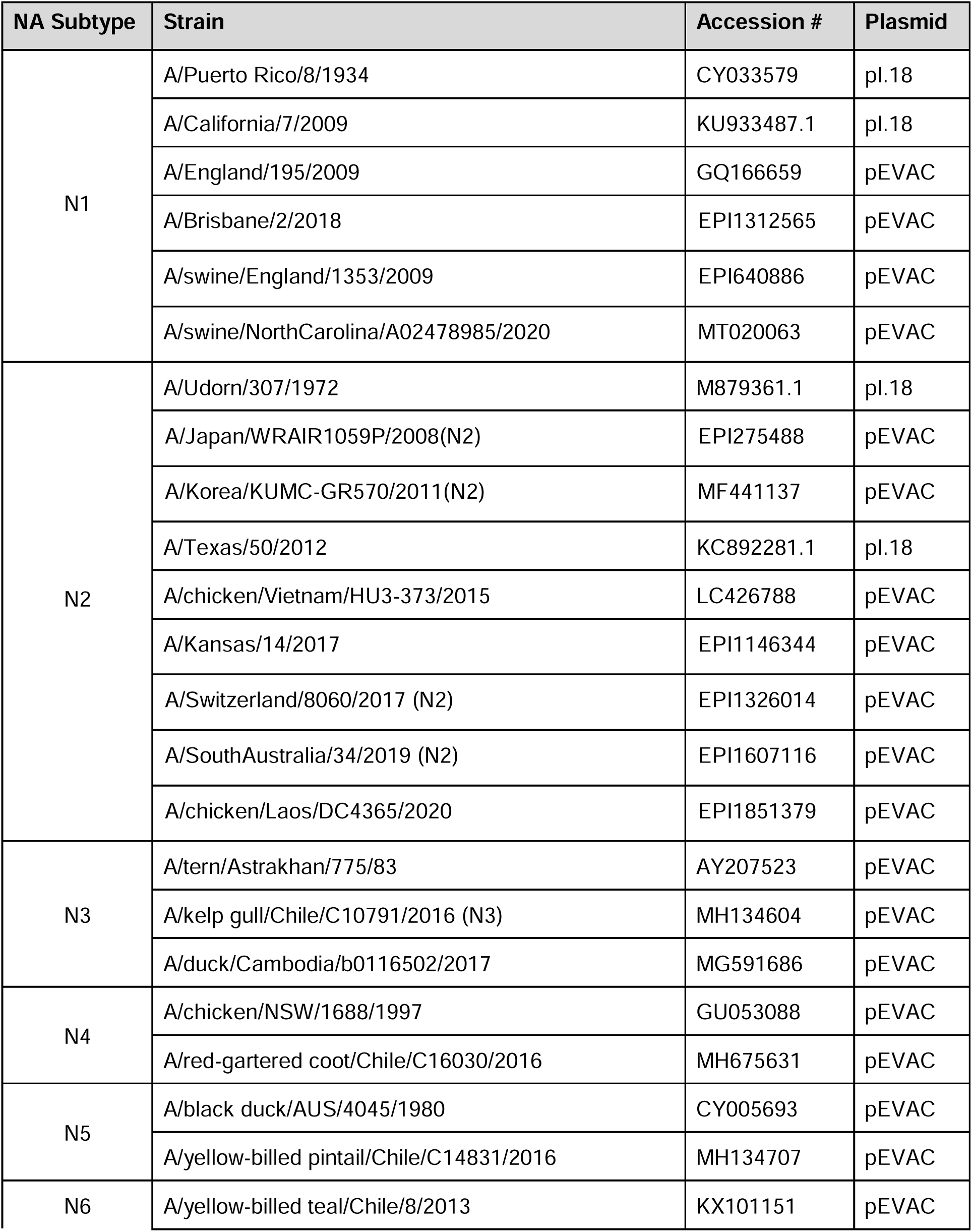
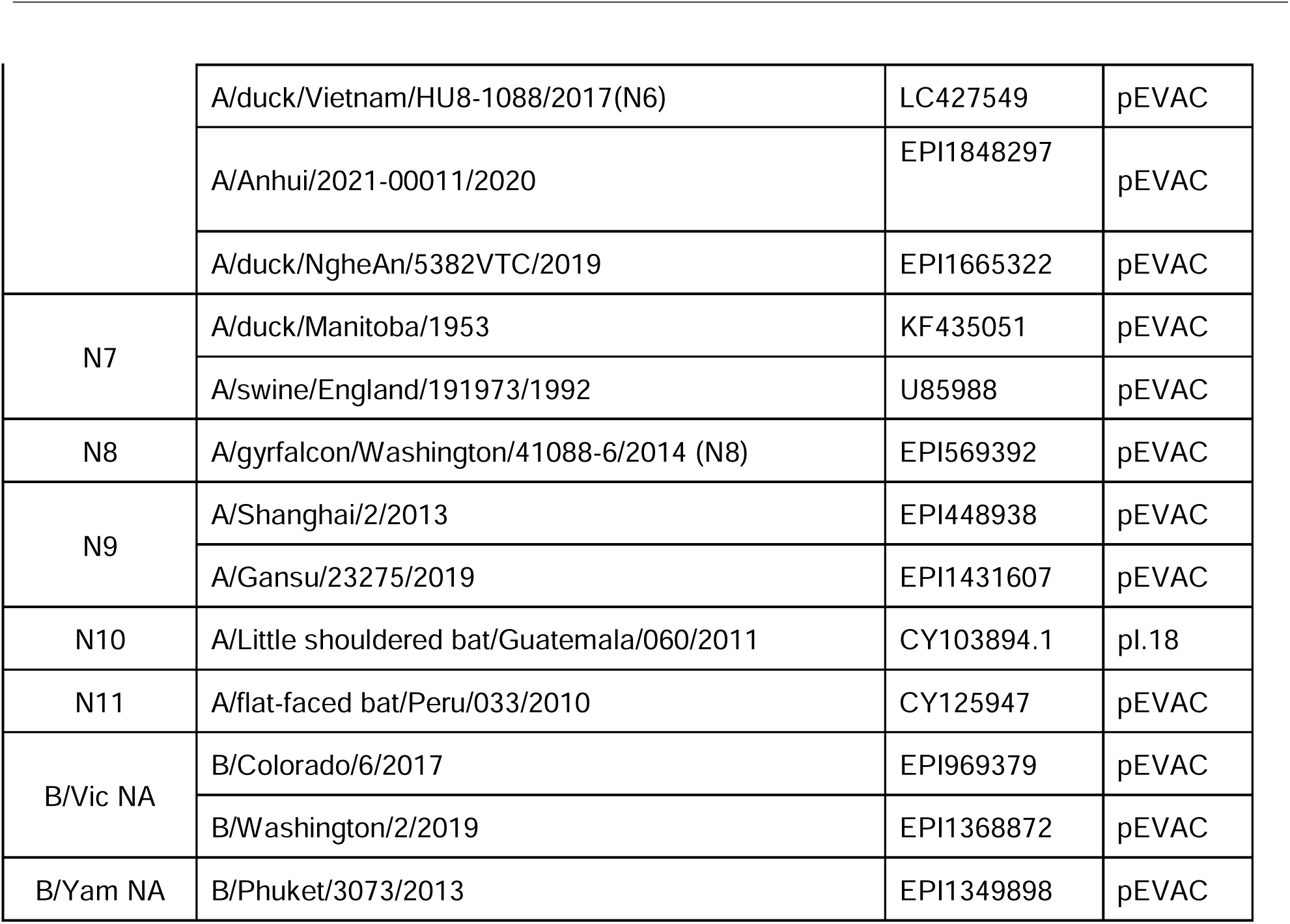
List of influenza neuraminidase pseudotypes (PV) available at the Viral Pseudotype Unit, University of Kent.

